## Supplement Materials for "Automated Classification of Bacterial Cell Sub-Populations with Convolutional Neural Networks"

Denis Tamiev, Paige Furman, Nigel Reuel

### Supplement 1: Bacterial strains, cell growth, and sample preparations.

*Bacterial Cell Sample Preparation:* B. subtilis 168 (NCBI reference sequence NC_000964.3) were used in this study. Cells were stored in glycerol stock format, picked, and cultured overnight in LB media. In the morning, the overnight was transferred into fresh media and grown till saturation. Cells fractions were harvested during early, logarithmic, and late stationary phases and prepared for imaging. Harvested cells were diluted or concentrated to absorbance at 600nm of 1 in 1ml culture tubes. A 1ml solution of cells was supplemented with 1ul of 10mg/ml solution of ethidium bromide. Cells were allowed to incubate with ethidium bromide at room temperature for 10min.

*Microscope Slide Preparation:* Imaging was performed on standard microscope slides (75 by 25mm). Adhesive rubber molds (Thermo fisher P24743) were applied to microscope slides. A 1% agarose/PBS solution was made, and casted in rubber molds. Molds were pressed with a microscope slide to have identical thicknesses of 0.5 mm.  A 1 ul solution of cells, incubated with ethidium bromide was applied to these agarose pads. Solution was allowed to sit at room temperature, protected from light, for 5min, to allow for liquid absorption. After that, agarose pads were covered with microscope cover slips to avoid drying of the gel.

*Image Acquisition:* All images were acquired on the Nikon Eclipse E800 microscope using FITC filter set. The images were acquired with CoolSnap HQ camera (Photometrics).The number of images per slide was limited to 25, to avoid photodegradation of ethidium bromide. In total, about 1000 microscope images was acquired and used in this experiment.

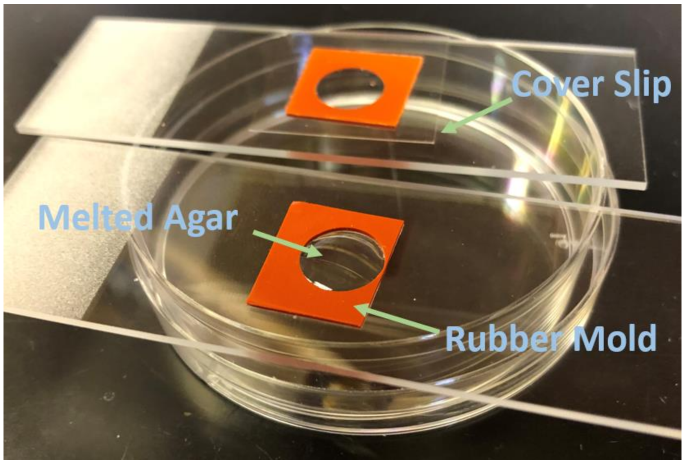

*Supplement Figure 1 – an example of agarose casting procedure (from ThermoFisher website).*

#### Supplement 2: Custom Algorithms for Normalizing Segment Size.

*Image Preprocessing:* All microscope images were first preprocessed with an adaptive binary function. Then, connected objects were segmented. Objects with total area of less than 20 square pixels were excluded. Coordinated of the bounding boxes of those segments were used to crop out portions of raw images. All segments were different in size, as they contained clusters of various numbers of cells. All segments had to be normalized in size prior to feeding into the CNN. Three approached to size normalizations were explored, to confine all segments to the 200 by 200 pixel dimensions (Figure 2).

*Various Size Normalization Algorithms:* In the null bumper approach, the segment was centered, and the distance between the edge of the cropped segment and the end of the normalized image was filled with pixels that were assigned null intensity. As a results, a frame of zero pixel intensities was created around all cropped images.

In the masked approach, the binary mask was used as a logical selection mask for pixels that belonged to the foreground (cell clusters) and background (agarose gel). The binary mask was applied to raw images, and cell clusters were extracted (Figure 2iv). The cluster was centered, and the background was filled with null intensity pixels. It was observed that the shape and morphology of cell clusters was significantly altered in this approach, and therefore, the networks were trained on masked images.

In the blended approach, the frame around the cell segment was blended with the background of the cell segment using a custom developed blending algorithm.

*Custom Image Blending Algorithm:* It was observed that in all segments, the most abundant pixel intensity belongs to the background, and the fluctuations of intensities of the background pixels is fairly narrow (around +/-10 pixels). To capitalize on that observation, the most popular pixel intensity of each segment was determined. Pixel intensities with values -7, -4, -3, -2, -1, 0, +1, +2, +3, +7 relative to the most popular pixel intensity were extracted. This matrix of pixels was called the seed matrix. The range of intensities around the most popular pixel intensity was determined empirically through trial and error. The selection criteria was based on how well blended images had appeared to the human operator.

It was observed that even the pixels from the background of the segment were never equal to zero. However, the bumper around the segment that was implemented to normalize all images by size contained only null intensity pixel. This provided a very simple selection criteria for determining where the frame of the image was located.

To blend the background of the segment with the null bumper frame, and make them appear as though they were continuous, pixel intensities from the seed matrix were randomly picked and assigned to the pixels that belonged to the frame. This allowed to achieve the desired result, and make the transition between the background of the cell segment, and the frame almost seamless.

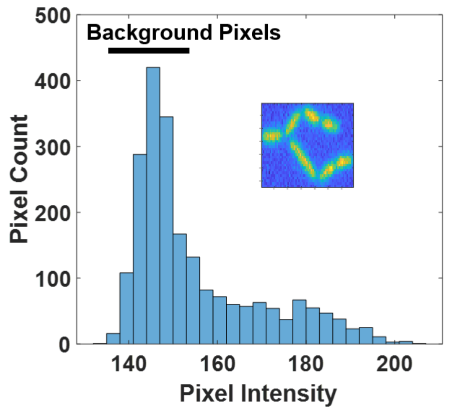

*Supplement Figure 2 – Histogram of pixel intensities in an image of a seven-cell cluster.*

All acquired cell segments were manually labeled to create a training, validation and testing image datasets (Supplement 13 – description of the custom labeling GUI). When all clusters from the acquired 1000 images were labeled, it was observed that, while instances of artifacts, and single cells were abundant, images with two or more cells in a cluster were significantly rarer (Figure 3). This presented a problem since training on the unbalanced dataset would result in an inaccurate network. To create a final image dataset, 81 images were randomly selected from each class of images. These 81 images were augmented and normalized to conform to a 200 by 200 pixel dimensions

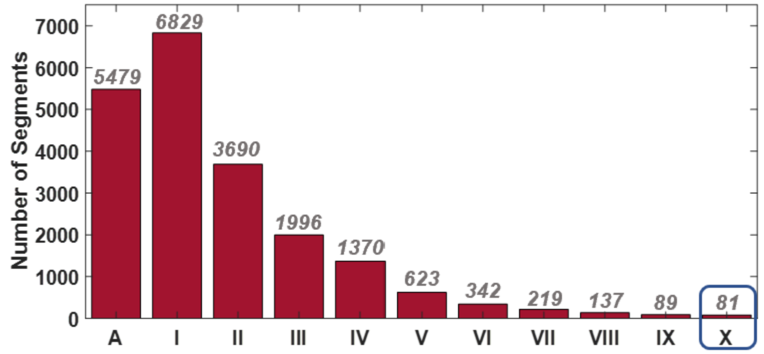

*Supplement Figure 3 – Histogram of all cluster types (A – artifact; I-X – single through ten cell clusters) extracted from raw microscope images. To normalize all data, 81 segment images were randomly selected of each cluster type.*

In order to increase the amount of data available for training, image datasets were augmented with rotation. For instance, the NB images were rotated at right angles (90 degrees) across both mirrored dimensions. Then, these images were placed in the center of a 200 by 200 array of pixel intensities (NB dataset). Alternatively, an additional step was taken (blending) to create the BB dataset. This resulted in an 8x increase in the amount of data that was available for training. Images were also rotated at fine angles (10 degrees), and across both mirrored dimensions. Then they were placed at the center of a 200 by 200 pixel array and blended with the background to create the AR dataset. This increased the amount of data available for training by 72 times.

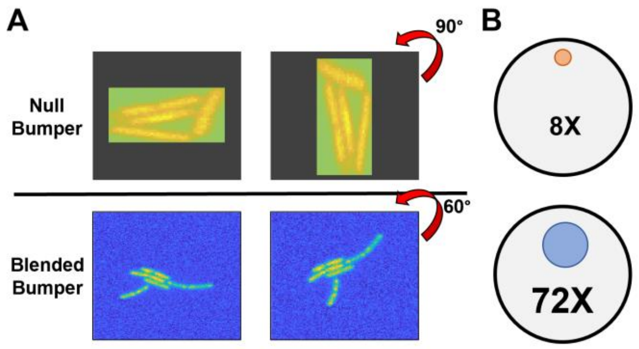

*Supplement Figure 4 – Panel "A" shows augmentation capacity with images that were normalized using the null bumper and blended bumper procedures. Panel "B" shows relative sizes of image datasets resulted from various augmentation procedures. Null bumper dataset was created by rotating inverted images by the right angles (8X). Advanced rotation dataset was created by rotating inverted images by 10-degree angles (72X).*

**Sample image preprocessing code implemented in Matlab:**

```
im=imread(thisImDir);
    %imagesc(im);

        %Extract image segments
        im2=imbinarize(im,'adaptive','Sensitivity', 0.55);

        rawBoxes=regionprops(im2,'BoundingBox');
        rawBoxes=reshape(struct2cell(rawBoxes),[length(rawBoxes) 1]);
        rawBoxes=cell2mat(rawBoxes);

        %% Extract area via regionprops (remove noise/single pixels)
        area=regionprops(im2,'Area');
        area=transpose({area.Area});
        area=cell2mat(area);

        rawBoxes(:,5)=area;

        %reshape rawBoxes array to remove small pixels
        selMat=rawBoxes(:,5)>10; %selection matrix (removes the noise based
on area size)
        a=rawBoxes(:,1);
        a=a(selMat);

        b=rawBoxes(:,2);
        b=b(selMat);

        c=rawBoxes(:,3);
        c=c(selMat);

        d=rawBoxes(:,4);
        d=d(selMat);

        rawBoxes=[];
        rawBoxes=[a, b, c, d];
```

```
        maxDX=max(rawBoxes(:,3));
        maxDY=max(rawBoxes(:,4));

        maxValues(i,1)=maxDX;
        maxValues(i,2)=maxDY;

        maxX=200;
        maxY=200;

        %create a dump folder for segments
            cd(thisDir);
            mkdir tempDump
            cd tempDump
            %reshape each segment

            if maxValues(i,1)<199 && maxValues(i,1)<199 %Excplude ims with
segments above input max
                 thisImName=int2str(i);

                    mkdir(thisImName);
                    cd(thisImName);
                for ii=1:size(rawBoxes(:,1))

                %get size of this box
                dX=rawBoxes(ii,3)-1; %shrunk by 1 to avoid going out of
bounds of the image due to rounding up.
                dY=rawBoxes(ii,4)-1; %same

                xBumper=int16(maxX-dX);
                xLeft=idivide(xBumper,2,'floor');
                xRight=maxX-dX-xLeft;

                yBumper=int16(maxY-dY);
                yTop=idivide(yBumper,2,'floor');
                yBottom=maxY-dY-yTop;

                %extract the image segment
                topLeftX=ceil(rawBoxes(ii,1));
                topLeftY=ceil(rawBoxes(ii,2));

                segIm=im(topLeftY:topLeftY+dY,topLeftX:topLeftX+dX);

                    % Expland the image
                    thisSegment=zeros(200,200);
                    thisSegment=uint16(thisSegment);
                    try
                    thisSegment(yTop:yTop+dY,xRight:xRight+dX)=segIm;
                    catch

                        %preds(:,:,i)=[];
                        break

                    end
                    %imagesc(thisSegment);
```

```
    %(!) Handover i and ii values;
    oldI=i;
    oldII=ii;
%Generate a randomized background matrix specific for each images.

h=histogram(segIm);

bins=h.BinCounts;
edges=h.BinEdges;

[val,idx]=max(bins);
seed=round(edges(idx));
blendMat=[seed-7,seed-4:1:seed+3,seed+7];

%Section above the image

blendSel=zeros(yTop-1,200);
for i=1:yTop-1
ranMat=randi(10,1,200);
for ii=1:200
blendSel(i,ii)=blendMat(ranMat(ii));
end
end

thisSegment(1:yTop-1,:)=blendSel; %!!!

%Section below the image
blendSel=zeros(200-yTop-dY,200);

for i=1:200-yTop-dY
ranMat=randi(10,1,200);
for ii=1:200
blendSel(i,ii)=blendMat(ranMat(ii));
end
end

thisSegment(yTop+dY+1:200,:)=blendSel;

%Section to the left of the image
blendSel=zeros(200,xRight-1);
for i=1:xRight-1
ranMat=randi(10,1,200);
for ii=1:200
blendSel(ii,i)=blendMat(ranMat(ii));
end
end

thisSegment(:,1:xRight-1)=blendSel;

%Section to the left of the image
blendSel=zeros(200,200-xRight-dX);
for i=1:200-xRight-dX
ranMat=randi(10,1,200);
for ii=1:200
blendSel(ii,i)=blendMat(ranMat(ii));
```

```
    end
    end

    thisSegment(:,xRight+dX+1:200)=blendSel;

    %imagesc(thisSegment);

    %(!) Handover old i and ii values to new ones
    i=oldI;
    ii=oldII;

preds(ii,:,i)=predict(net,thisSegment);
thisPreds=preds(ii,:,i);
[M1,I1]=max(thisPreds);
thisPreds(I1)=0;
[M2,I2]=max(thisPreds);
thisPreds(I2)=0;
[M3,I3]=max(thisPreds);
```

#### Supplement 3: General Workflow.

The conventional workflow of selecting, training, and deploying neural networks is a well-established process[1–3]. First, training and evaluation data needs to be acquired. For supervised ML these data sets are then manually labeled as depicted on the Supplement figure 5A (Supplement 13). As discussed in the main manuscript, the quality of preprocessed data determines the accuracy of the network, and, as such, special considerations should be given to image preprocessing. Often, a general preprocessing algorithm is chosen, and then, during the fine tuning of the network, other algorithms are explored. For example, the raw image can be reduced in resolution to match the input layer of the pre-trained network during the first attempt at training the network. Depending on the performance of the network the user can try reducing the size of the raw image to a variety of resolutions and train networks on those images, until the optimal performance of the network is achieved.

In cases where data sets are more sparse (e.g. microscope images), various data augmentation mechanisms such as image rotation, addition of random noise and many others can be implemented to decrease the amount of labeled images needed to train a network.

Then, an appropriate network architecture needs to be either designed or selected. The most common approach is to begin with picking a handful of pre-trained networks, and perform transfer learning. The network can be trained either on a local resource (a desktop computer with a powerful GPU), or on a cloud resource (AWS, Microsoft Azure, Google Cloud, etc.) as depicted in the Supplement Figure 5B.

In the next phase, trained networks are evaluated with reserved data that they have not encountered during the training process (Supplement Figure 5C). This can also be done either on a local (desktop with GPU) or remote (cloud service) resources. Best-performing networks, are then selected for fine-tuning. This can be achieved by (1) testing various image preprocessing algorithms, (2) providing more data to train on, (3) optimizing training conditions (learning rate, number of iterations, etc). Typically, fine-tuning a network requires a lot of empirical experience, and a deeper understanding of network's structures as well as the training mechanisms. It is best to follow the suggestions made in the paper that originally describes the design of the pre-trained network that was chosen.

When a satisfactory accuracy is achieved, the network is deployed on new application data. In our case, the network was deployed as a classifier of cell clusters for microscope images. The output of the network was the cell count within a cluster, and the total count was the numerical sum of the individual outputs (Supplement figure 5D).

In this paper, we used the cCNN that was demonstrated to work well with images that contained objects with similar features (circular clusters of bacterial cell colonies on petri dishes) to what we observed on images used in this study.

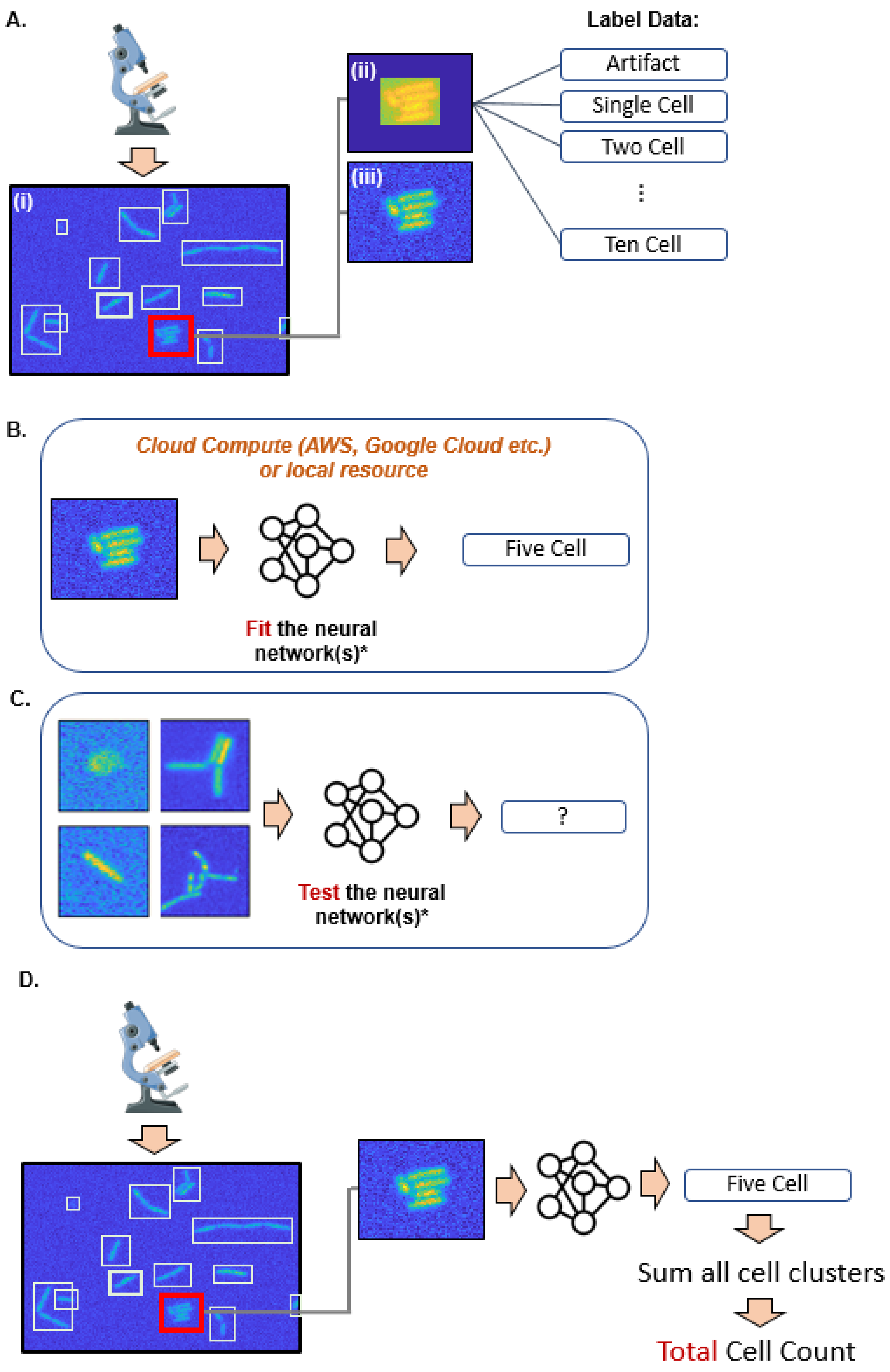

*Supplement Figure 5 – Conventional workflow for selecting, training and deploying neural networks. (A) Dada preprocessing. Acquire microscope images, and segment into clusters (i). Normalize all data (ii & iii). Label images. (B) Select a neural network*

*or pick several networks for comparison. Train ("fit") network with normalized images and labels. (C). Evaluate the accuracy of the network, and compare various network architectures. (D). Deploy the solution.*

### Supplement 4: CNN Architecture.

In this paper a classification Convolutional Neural Network (cCNN) was employed. The architecture of the network was adapted from *Ferarri et al* with few notable modifications (Supplement Figure 6) [4]. The input layer was modified to accept 200 by 200 pixel images of cell segments. The output layer was modified to classify images in one of 11 classes – artifacts, single cells, and two through ten cell clusters. In this paper, the network was trained on one of the three image datasets – null bumper, blended bumper, advanced rotation. Consecutively, this resulted in generation of 3 different cCNNs.

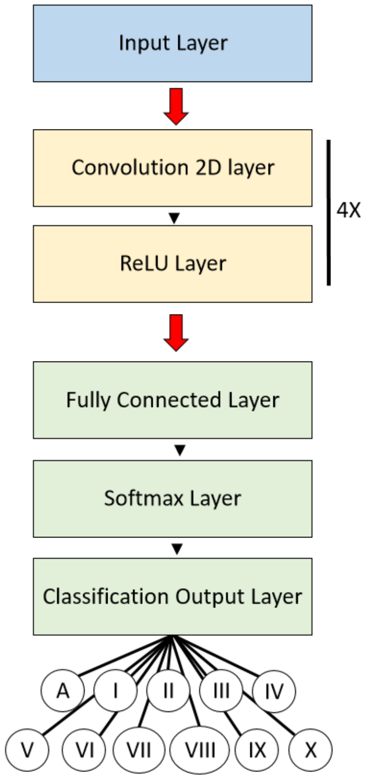

*Supplement Figure 6 – Graphical representation of the architecture of the CNN used in this project. Input layer is highlighted in blue, middle layers (4 repeats) are highlighted in yellow, and output layers are highlighted in green.*

### Supplement 5: Training Conditions.

All programmatic procedures, including training and implementation of cCNN was performed in Matlab (version 2019a). Matlab was implemented in Amazon Web Services cloud computing environment. Specifically, EC2 cluster p2.xlarge was used (NVIDIA K80 GPU).

The cloud environment was configured with a AWS CloudFormation template that was preconfigured for Matlab as described elsewhere[5]. Microscope images were transferred to and from the cloud service via the secure copy protocol (SCP), while larger files such as trained networks were transferred via Amazon's S3 service.

Training of the null bumper cCNN was performed for 10 Epochs with validation. Initial training rate was set to $1^{-4}$, and was programmed to decrease in by 10 times every epoch after the third Epoch. It was observed that after the third epoch the validation and training accuracies as well as loss, have stabilized. The initial training rate was scheduled to decrease after it stabilized in order to fine-tune the network's fit. Training was performed with stochastic gradient descent, and batch size of 20 images was selected. First applying momentum was set to 0.9.

Training of blended bumper and advanced rotation datasets was stopped as soon as the accuracy of the validation set was surpassed by the training set.

Null bumper and blended image datasets were parsed to dedicate roughly 85% of images to training 5% to validation and 10% to testing. Advanced rotation dataset was parsed to dedicate 90% of images to training and 5% to validation and to testing.

Training on the null bumper dataset (NB) did not result in overfitting of the data during the first five epochs. The network was underfitting, as evident by plateaued validation accuracy. Each epoch consisted of 282 iterations. This is evident from the training accuracy data fluctuating around a similar value as the validation accuracy (Supplement Figure 7). While the amplitude of fluctuations increased during later epochs, even after ten epochs the training accuracy had stabilized at around 60% (data not shown). This signified that the training capacity was reached prior to overfitting.

In comparison, when the same network was trained on a blended bumper dataset, significant overfitting began immediately after the first epoch. This indicates that the learning capacity of this network is sufficient to train a highly accurate solution when microscope images are preprocessed using our custom algorithm. To eliminate the plateaued accuracy of the network on the validation set, we generated more training data through augmentation (AR dataset), and trained a third network.

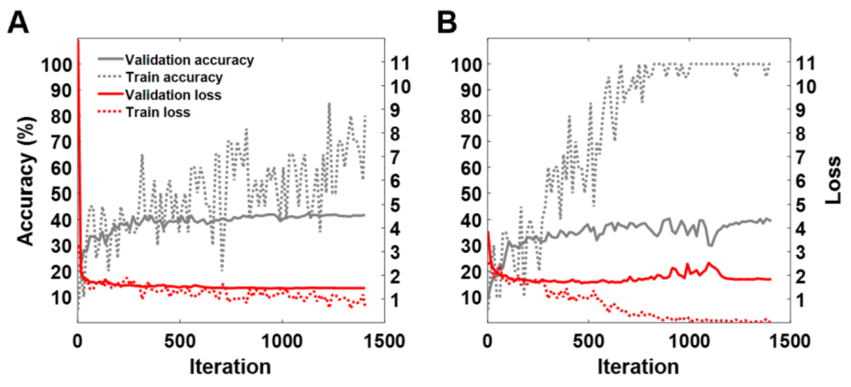

*Supplement Figure 7 – Training progress during first 5 epochs (282 iterations per epoch) of networks trained on (A) null, and (B) blended bumper datasets.*

#### ***Example of the code used on the AWS cluster:***

```
function transferLearning_V2_EC2
clc
clear
close all

%% Section 1: generate a network from scratch

% Initiate all layers here
inLayer = imageInputLayer([150 150]);
convLayer1 = convolution2dLayer([5 5], 20);
convLayer2 = convolution2dLayer([5 5], 50);
convLayer3 = convolution2dLayer([5 5], 100);
convLayer4 = convolution2dLayer([5 5], 200);

midLayers = [reluLayer()];
outLayers = [fullyConnectedLayer(4); softmaxLayer(); classificationLayer()];

layers = [inLayer; convLayer1; midLayers; convLayer2; midLayers; ...
     convLayer3; midLayers; convLayer4; midLayers; outLayers];

%% Section 2: Create Image Datastores (comment out after initiation)

%load table from the directory
thisDir=pwd;
%load(string([thisDir, '/augmented_v2', '/selected55/', 'augmentedSelected.mat'])); %(!) CHANGE
'\' to '/'
load('C:\Users\Denis Tamiev\Google Drive\190310 Den Main folder\190310 Neural Network for spore
detection\software\Version 2\Project 1 (Fluorescent images of vegetative
cells)\Images\database\augmented_v2\selected55\augmentedSelected.mat');

%mutate source to reflect current directory.
imdsDir=strrep(augmentedSegments.segment,"C:\Users\ddtamiev\Google Drive\190310 Den Main
folder\190310 Neural Network for spore detection\software\Version 2\Project 1 (Fluorescent images
of vegetative cells)\Images\database", thisDir);
imdsDir=strrep(imdsDir,'\','/');
augmentedSegments.ClusterType;
h=histogram(augmentedSegments.ClusterType);
binCs=h.BinCounts;
imds=imageDatastore(imdsDir,'Labels', categorical(augmentedSegments.ClusterType));

h2=histogram(imds.Labels);
binCs2=h2.BinCounts;

[trainds,valds,testds] = splitEachLabel(imds,0.8,0.1,'randomized');
```

```
%save these datastores for additional training.
filename=char([thisDir, '/datastores/trainds']);
save(filename,'trainds')
filename=char([thisDir, '/datastores/trestds']);
save(filename,'testds')
filename=char([thisDir, '/datastores/valds']);
save(filename,'valds')

%% Section 3: Training options
options = trainingOptions('sgdm','MaxEpochs',10,'InitialLearnRate',0.0001,'Plots','training-progress','ValidationData', valds, ...
    'ValidationFrequency', 10, 'ValidationPatience', 5, 'LearnRateSchedule', 'piecewise', ...
    'LearnRateDropFactor', 0.001, 'LearnRateDropPeriod', 3,'ExecutionEnvironment', 'gpu', 'OutputFcn',@info);

%% Section 4: Train the network

net=trainNetwork(trainds,layers,options);
filename=char([thisDir, '/outputVars/outputAll']);
save(filename);
end
```

#### Supplement 6: Activations from Convolution Layers.

During the original stages of the project, we first trained cCNN on the null bumper image dataset. We have observed that the accuracy of the network reached its limit at around 45% (for the validation dataset). We were surprised to observe that the network was never able to overfit the training data, even when the learning rate was not scheduled to decrease. Next, we decided to explore various pathways to increasing the networks accuracy beyond 45%. We first investigated the activations from all convolution layers (Supplement Table 1). It was observed that the null bumper network was focusing on the frame of image segments rather than on the shape of cell clusters. This was especially predominant towards the latter convolution layers. As a result, by the time the image makes its way to the last convolution layer, the network was no longer able to recognize any of the cells inside of the image.

To address this issue we decided to explore various ways to remove the rigid boundary between the image segment and the null bumper. This was achieved by developing a blending algorithm which was described earlier in this supplement section. As a result, we developed a blended image dataset. All other training conditions were otherwise kept identical to the null bumper scenario.

Almost instantly (after the first epoch) we observed significant overfitting. By the thirds epoch, accuracy of the network tested on the training image dataset reached 100%, while it stabilized at around the same value (about 45%) for the validation dataset.

We tried to decrease the learning rate of the network, but only a slight increase in accuracy (tested on the validation set) was observed. Therefore, we decided to explore strategies for increasing the size of our training dataset through augmentation to improve the accuracy of our cCNN.

Image augmentation was restricted to rotation, and inversion algorithms. Through rotating segments of cell clusters by 10 degrees over to mirror conformations, a significant increase in the size of the database was achieved (72X). When the network was trained on this increased larger image dataset, about a 14% increase in accuracy was achieved.

*Supplement Table 1 – Activations of Convolution Layers of cCNN trained on either blended bumper or null bumper image datasets.*

| **Null bumper cCNN activations** | **Blended bumper cCNN activations** |
|---|---|
| 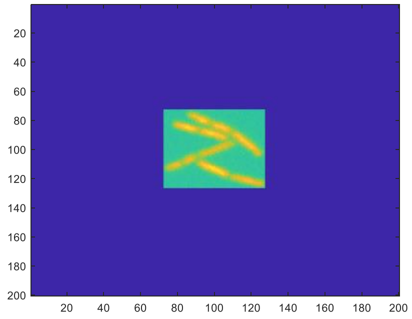 | 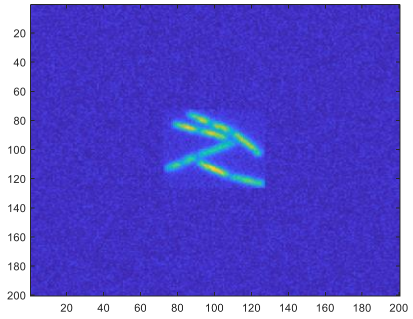 |
| *Convolution Layer 1* 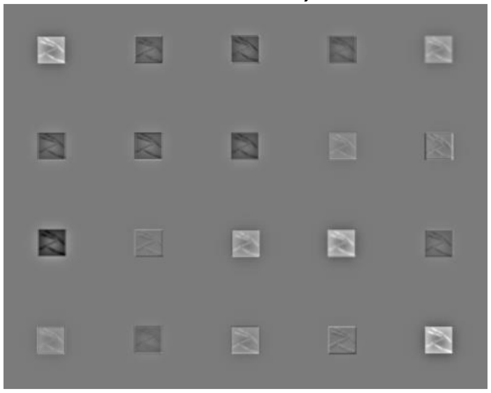 | *Convolution Layer 1* 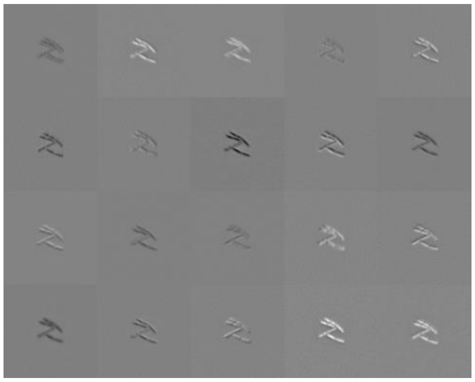 |
| *Convolution Layer 2* 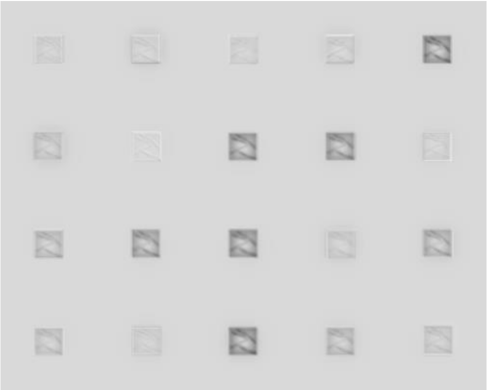 | *Convolution Layer 2* 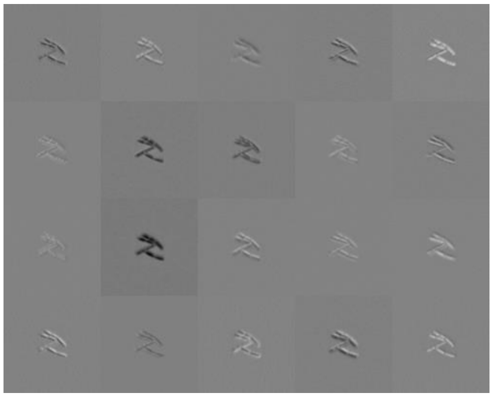 |
| *Convolution Layer 3* | *Convolution Layer 3* |

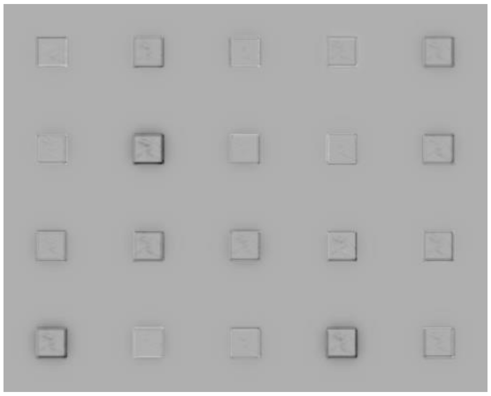

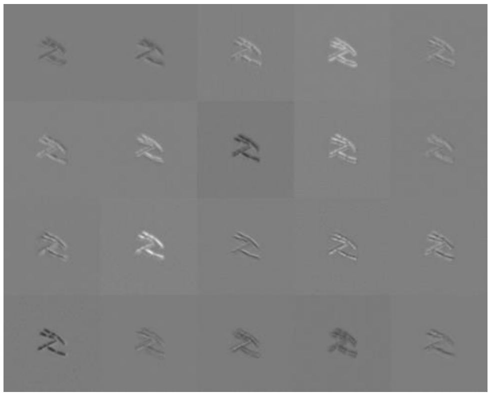

*Convolution Layer 4*

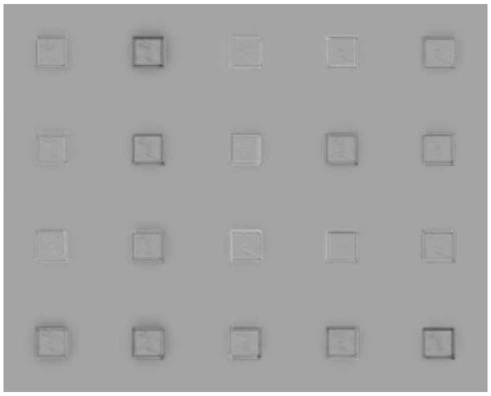

*Convolution Layer 4*

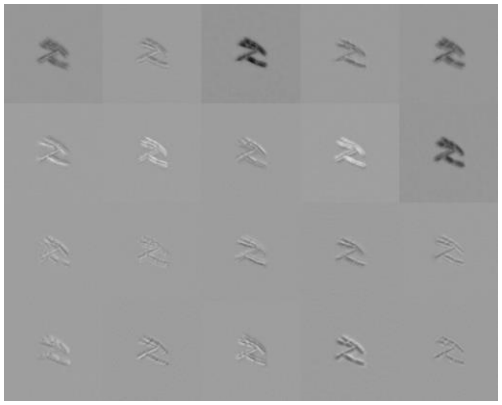

#### Supplement 7: Confusion Matrixes of Networks’ Accuracies.

After training of the networks was complete, they were tested on both validation, and testing datasets, to evaluate their performance, capacity to effectively classify cell clusters. Accuracy data was represented in the confusion matrix format (Supplement Figure 8).

Correct predictions are highlighted in grey and lay along the diagonal. This data can be reduced to the bar chart format by plotting correct predictions (number on the diagonal) in a bar chart for each class (for every row). This data was presented in the main manuscript.

We observed that there is spatial relationship in between the classes presented in the confusion matrix. Specifically, a cluster of 5 cells is more similar to the cluster of 4 and 6 cells. Therefore, a new metric was designed to evaluate the accuracy not only by the number of exact guesses but also by how far the subpar guesses are located from the diagonal. The rules for designing this metric is described in detail in Supplement 8.

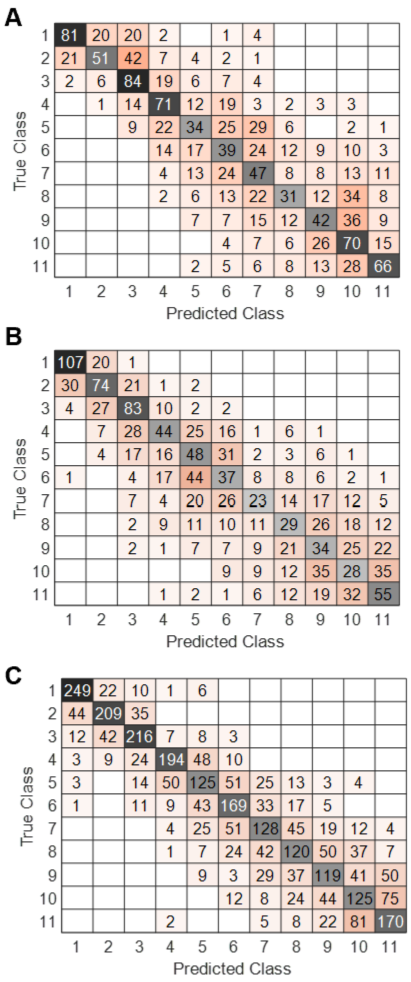

*Supplement Figure 8 – Confusion Matrix representation of the accuracy data acquired from testing networks trained on (A) null bumper, (B) blended bumper, (C) advanced rotation datasets.*

### Supplement 8: Accuracy Data extracted from Confusion Matrixes.

As mentioned in Supplement 7, correct guesses (numbers along the diagonal) were presented in the bar chart in the main manuscript. The exact values used for the bar chart could be found in the supplement table 2.

*Supplement Table 2 – unweighted accuracy values extracted from the confusion matrix.*

| | unweighted accuracy | | |
|---|---|---|---|
| | **Null Bumper** | **Blended Bumper** | **Advance Rotation** |
| **A** | 0.63281 | 0.83594 | 0.86458 |
| **I** | 0.39844 | 0.57813 | 0.72569 |
| **II** | 0.65625 | 0.64844 | 0.75 |
| **III** | 0.55469 | 0.34375 | 0.67361 |
| **IV** | 0.26563 | 0.375 | 0.43403 |
| **V** | 0.30469 | 0.28906 | 0.58681 |
| **VI** | 0.36719 | 0.17969 | 0.44444 |
| **VII** | 0.24219 | 0.22656 | 0.41667 |
| **VIII** | 0.32813 | 0.26563 | 0.41319 |
| **IX** | 0.54688 | 0.21875 | 0.43403 |
| **X** | 0.51563 | 0.42969 | 0.59028 |
| **Average** | 0.4375 | 0.39915 | 0.57576 |

#### Supplement 9: Confidence of the Network by Class.

In this project we investigated the confidence values of predictions for each output class. As mentioned above, to test the accuracy of the null bumper, and blended bumper networks 128 images of each class were used (Advanced rotation network was tested with 228 images from each class). Then, the confidence values for each class were calculated and presented in a histogram format. As a result, three sets of 11 histograms (one for each network), one for each class, were created. Average confidence of each class was then extracted, for each of the networks, resulting in 33 values (11 classes, and 3 networks). These values were used to generate the radar plot in figure 8D of the main manuscript. Individual values are presented in Supplement Table 3, and individual histograms are presented in Supplement Figure 9.

*Supplement Table 3 – Average confidence values for each output class.*

| | Null Bumper | Random Bumper | Advanced Rotation |
|---|---|---|---|
| **Artifact** | 0.7482 | 0.82526 | 0.848176 |
| **One** | 0.54473 | 0.73333 | 0.765167 |
| **Two** | 0.53571 | 0.65643 | 0.685428 |
| **Three** | 0.48721 | 0.59556 | 0.687486 |
| **Four** | 0.39304 | 0.53293 | 0.505335 |
| **Five** | 0.37868 | 0.5587 | 0.488008 |
| **Six** | 0.37022 | 0.49299 | 0.47094 |
| **Seven** | 0.37254 | 0.52955 | 0.444413 |

| | | | |
|---|---|---|---|
| **Eight** | 0.46331 | 0.63501 | 0.528326 |
| **Nine** | 0.49652 | 0.61563 | 0.531822 |
| **Ten** | 0.60012 | 0.61033 | 0.667382 |
| **Average** | 0.49002 | 0.61688 | 0.602044 |

*Supplement Figure 9 – Average confidence values for each output class presented in the histogram format.*

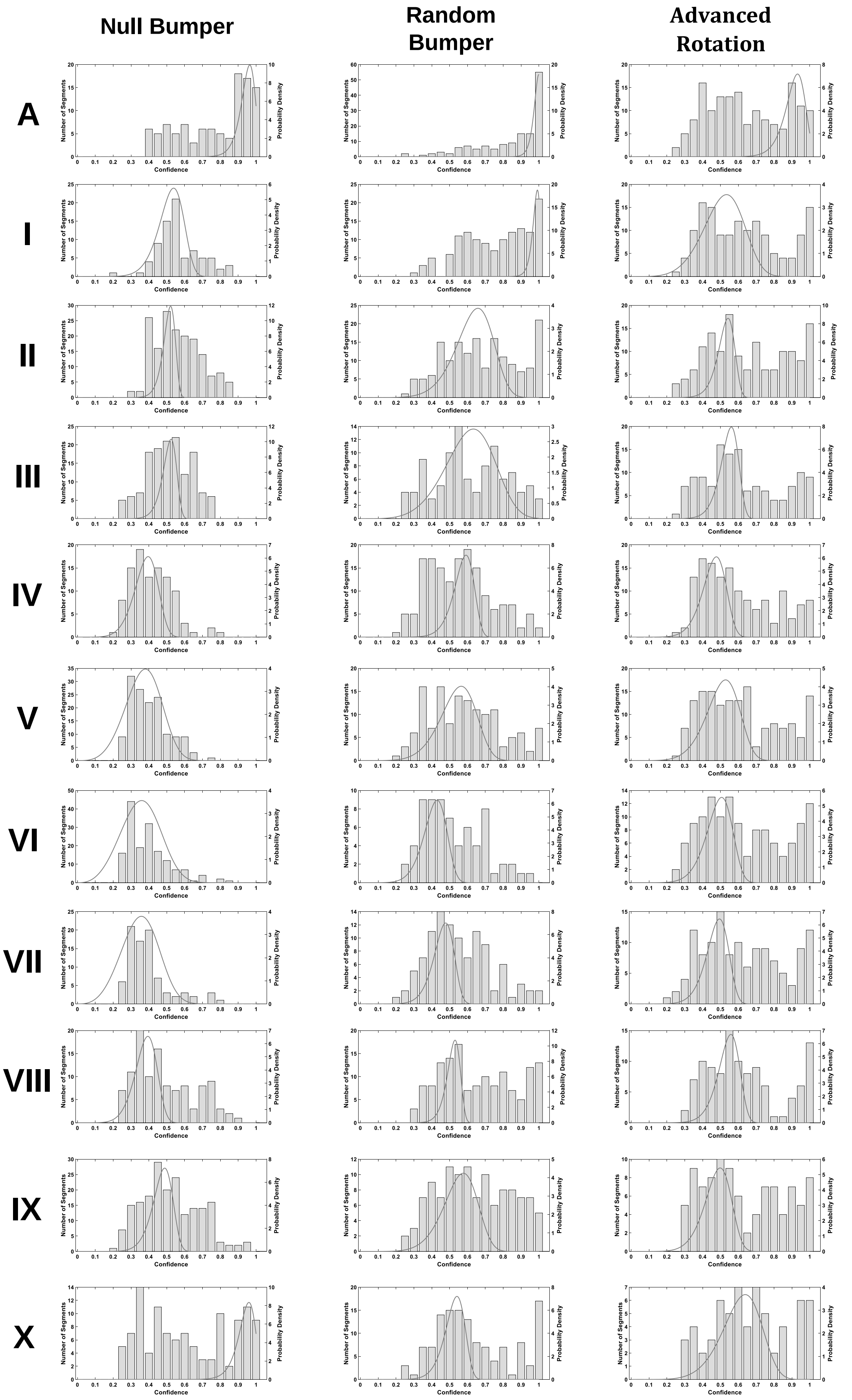

#### Supplement 10: Protocol for Manual Cell Counting.

In figure 11 of the main manuscript we show how various counting methods, 3 different cCNNs and ImageJ, compare to manual counting. Manual counting was performed by overlaying gridlines over the image in PowerPoint, printing that image, and manually counting individual cells with a clicker. In this case, all cell counts were performed by one person. All images were counted three times, by the same person, and the average count was reported in the Supplement Table 4.

Raw data that was used to in the scatter plots (figure 11 of the main manuscript) is presented in the supplement table 4.

*Supplement Table 4– Raw data used for the scatter plot in figure 11 of the main manuscript.*

| im# | Null bumper | Blended bumper | Epochs | | | | ImageJ | Human |
|---|---|---|---|---|---|---|---|---|
| | | | Adv. Rotation 1 | Adv. Rotation 2 | Adv. Rotation 3 | Adv. Rotation 4 | | |
| 1 | 33 | 14 | 16 | 16 | 16 | 15 | 32 | 21 |
| 2 | 46 | 27 | 30 | 27 | 28 | 27 | 38 | 32 |
| 3 | 28 | 14 | 12 | 14 | 15 | 12 | 18 | 13 |
| 4 | 60 | 39 | 39 | 36 | 36 | 33 | 45 | 28 |
| 5 | 2 | 1 | 2 | 2 | 2 | 1 | 1 | 1 |
| 6 | 16 | 15 | 11 | 11 | 11 | 11 | 19 | 10 |
| 7 | 21 | 12 | 16 | 18 | 18 | 18 | 16 | 14 |
| 8 | 28 | 22 | 23 | 22 | 22 | 22 | 22 | 24 |
| 9 | 10 | 7 | 5 | 6 | 6 | 6 | 8 | 7 |
| 10 | 17 | 14 | 13 | 13 | 13 | 12 | 17 | 18 |
| 11 | 132 | 100 | 91 | 90 | 90 | 88 | 84 | 98 |
| 12 | 94 | 71 | 59 | 57 | 56 | 56 | 68 | 58 |
| 13 | 199 | 170 | 173 | 168 | 168 | 157 | 132 | 161 |
| 14 | 97 | 73 | 73 | 68 | 68 | 67 | 64 | 75 |
| 15 | 119 | 92 | 84 | 95 | 95 | 86 | 68 | 79 |
| 16 | 131 | 109 | 103 | 105 | 111 | 105 | 88 | 109 |
| 17 | 161 | 129 | 131 | 136 | 130 | 131 | 105 | 131 |
| 18 | 103 | 76 | 81 | 73 | 73 | 72 | 84 | 43 |
| 19 | 58 | 41 | 39 | 38 | 38 | 35 | 40 | 43 |
| 20 | 62 | 46 | 40 | 42 | 40 | 43 | 43 | 51 |
| 21 | 68 | 54 | 49 | 51 | 51 | 51 | 69 | 57 |
| 22 | 67 | 51 | 51 | 47 | 46 | 47 | 54 | 52 |
| 23 | 248 | 187 | 180 | 194 | 192 | 166 | 145 | 174 |
| 24 | 98 | 79 | 74 | 71 | 71 | 67 | 68 | 70 |

### Supplement 11: Watershed Method of Image Segmentation Implemented in ImageJ

ImageJ is a common free image analysis package initial developed at the NIH to support microscopy and radiology work[6]. One use of ImageJ is to segment microscope images into clusters of cells using binary thresholding, similarly to how such clusters were first identified using Matlab in this paper. To count cells, ImageJ can then employ a watershed algorithm (treating pixel intensity as topology) to parse those cell clusters into individual cells. Since its original development, various modifications of the classical watershed algorithm were improved[7]. Additional customization for a specific application is possible with a goal to minimize either over- or under-segmentation. ImageJ software implements one version of this algorithm, and, without further customization, our images were under segmented (Supplement figure 10).

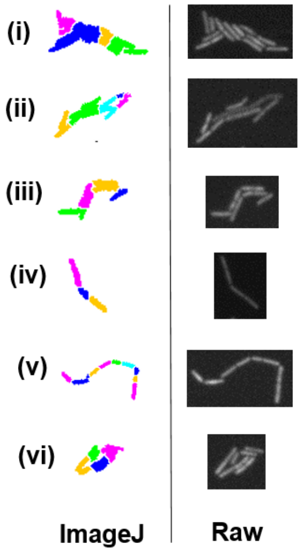

*Supplement Figure 10 – representative images of cell clusters processed in ImageJ (watershed), and corresponding raw images (i, ii, iii, vi – "side-to-side" clusters; iv, v – "end-to-end" clusters).*

Consequently, the performance of the ImageJ approach that relies on selecting cells based on size and parsing cell clusters using a watershed algorithm varies depending on the application. When working with images that depict a few multicellular clusters, ImageJ can be used successfully. Images with high concentrations, including biofilms, are less appropriate for this approach.

### Supplement 12: Human Performance Experiment.

*<u>Human Performance Experiment</u>* – a human performance study was created under the IRB's oversight (IRB ID 19-566). The study was classified as an exempt.

**<u>Subject selection:</u>**

In this study 5 test subjects were recruited. Test subjects were selected to be both men and women in the 21-30 year old age range. Subject's identities were not recorder to protect their identities.

**<u>Testing conditions:</u>**

Test subjects were given 25 images acquired with a fluorescent microscope. Out of those 25 images, 15 were unique and 5 were repeated in random order twice (an example image used for this study can be found at the end of this section. All images used for this study will be made available upon a request submitted to the corresponding author). Images contained square grids placed after the fact to mimic appearance of a cytometer microscope slide. All images were pasted in the power points slide and were presented to test subjects. Test subjects were placed in a conference room with a supervisor who was recording their processing speed. The subjects were counting cells from a PowerPoint slide presented to them on a computer screen, using a cell counter. Test subjects then self-reported the number of cells that they counted on each image.

**<u>Data processing:</u>**

Two types of data were recorder – (1) how long it took each subject to count cells on each microscope image, and (2) how many cells test subjects counted on each image. The average cell count per image between 5 test subjects was calculated to extract the standard deviation. Then, the standard deviation for each image was plotted as a bar chart (Figure 6A, main manuscript).

Data acquired from a cCNN based computer algorithm was acquired by running the algorithm on the same 25 images, and the cell count as well as the processing time were recorded. The algorithm was applied to 25 images 5 times, mimicking 5 separate human test subjects. Detailed description of the algorithm can be found below (Figure 6A, main manuscript).

Average standard deviation was calculated by taking an average of individual 25 standard deviation values for both human test subjects and the algorithm (Figure 6A, inset, main manuscript).

Individual cell counts and processing time for test subjects are recorded in Supplement Tables 6, and 7 for the cCNN based algorithm. Cells count on repeat images was recompiled from Supplement Table 5 into Table 8 for the ease of comprehension (Figure 6B, main manuscript).

*Supplement Table 5 – Processing time and cell count acquired from test subjects during the human performance experiment.*

| Image # | Processing time (seconds) Person 1 | Person 2 | Person 3 | Person 4 | Person 5 | Cell Count per image (cells) Person 1 | Person 2 | Person 3 | Person 4 | Person 5 | STDEV |
|---|---|---|---|---|---|---|---|---|---|---|---|
| 1 | 15 | 21 | 22 | 6 | 8 | 23 | 24 | 16 | 19 | 16 | 3.78 |
| 2 | 25 | 30 | 23 | 11 | 12 | 37 | 35 | 21 | 26 | 21 | 7.62 |
| 18 | 92 | 68 | 60 | 53 | 62 | 164 | 145 | 89 | 142 | 95 | 33.11 |
| 56 | 99 | 72 | 59 | 53 | 70 | 194 | 175 | 102 | 143 | 112 | 39.51 |
| 4 | 18 | 26 | 17 | 14 | 15 | 36 | 32 | 22 | 27 | 22 | 6.18 |
| 6 | 7 | 11 | 4 | 4 | 4 | 13 | 11 | 4 | 8 | 6 | 3.65 |
| 2 | 19 | 21 | 11 | 14 | 10 | 35 | 32 | 19 | 25 | 21 | 6.91 |
| 56 | 87 | 68 | 47 | 59 | 66 | 192 | 173 | 106 | 136 | 113 | 37.46 |
| 1 | 17 | 19 | 10 | 10 | 8 | 28 | 23 | 15 | 20 | 16 | 5.32 |
| 8 | 13 | 17 | 7 | 7 | 6 | 27 | 24 | 11 | 18 | 13 | 6.88 |
| 15 | 42 | 40 | 26 | 30 | 34 | 94 | 81 | 48 | 67 | 54 | 18.97 |
| 17 | 33 | 33 | 23 | 26 | 24 | 67 | 58 | 41 | 53 | 44 | 10.55 |
| 21 | 52 | 50 | 28 | 35 | 32 | 99 | 110 | 53 | 81 | 54 | 25.81 |
| 18 | 65 | 58 | 40 | 42 | 53 | 151 | 149 | 85 | 125 | 91 | 31.20 |
| 21 | 52 | 52 | 29 | 31 | 42 | 98 | 103 | 56 | 91 | 62 | 21.53 |
| 19 | 35 | 38 | 20 | 29 | 28 | 68 | 74 | 40 | 61 | 42 | 15.33 |
| 20 | 34 | 32 | 26 | 23 | 28 | 70 | 73 | 48 | 53 | 55 | 11.03 |
| 22 | 52 | 46 | 28 | 40 | 40 | 119 | 119 | 64 | 94 | 71 | 25.87 |
| 29 | 41 | 29 | 27 | 24 | 27 | 77 | 77 | 50 | 64 | 56 | 12.19 |
| 51 | 19 | 24 | 12 | 14 | 11 | 43 | 45 | 20 | 34 | 23 | 11.34 |
| 52 | 9 | 25 | 13 | 13 | 13 | 46 | 49 | 23 | 33 | 27 | 11.48 |
| 54 | 24 | 38 | 12 | 15 | 18 | 58 | 55 | 29 | 45 | 29 | 13.83 |
| 55 | 20 | 21 | 11 | 14 | 12 | 48 | 47 | 27 | 42 | 28 | 10.21 |
| 58 | 30 | 22 | 16 | 21 | 24 | 69 | 68 | 43 | 59 | 51 | 11.14 |
| 59 | 23 | 30 | 11 | 19 | 19 | 55 | 54 | 29 | 41 | 33 | 11.87 |
| TOTAL: | **923** | **891** | **582** | **607** | **666** | | | | | **AVG** | **15.71** |
| | | | | | | | | | | **MAX** | **39.51** |
| | | | | | | | | | | **MIN** | **3.65** |

*Supplement Table 6 – Processing time and cell count acquired from the cCNN based algorithm during the human performance experiment.*

| Image # | Processing time (seconds) 1st | Cell Count per image (cells) 1st | 2nd | 3rd | 4th | 5th | STDEV |
|---|---|---|---|---|---|---|---|
| 1 | 4.65 | 14 | 19 | 16 | 17 | 15 | 1.92 |
| 2 | 4.39 | 27 | 28 | 27 | 25 | 27 | 1.10 |
| 18 | 17.75 | 170 | 172 | 173 | 171 | 173 | 1.30 |
| 56 | 21.62 | 187 | 181 | 179 | 178 | 184 | 3.70 |

| | | | | | | | | | | | |
|---|---|---|---|---|---|---|---|---|---|---|---|
| 4 | | | | | 6.40 | 39 | 44 | 41 | 37 | 44 | 3.08 |
| 6 | | | | | 2.13 | 15 | 15 | 15 | 15 | 15 | 0.00 |
| 2 | | | | | 4.39 | 27 | 28 | 27 | 25 | 27 | 1.10 |
| 56 | | | | | 21.62 | 187 | 181 | 179 | 178 | 184 | 3.70 |
| 1 | | | | | 4.65 | 14 | 19 | 16 | 17 | 15 | 1.92 |
| 8 | | | | | 2.78 | 22 | 23 | 22 | 23 | 22 | 0.55 |
| 15 | | | | | 10.60 | 100 | 100 | 102 | 98 | 99 | 1.48 |
| 17 | | | | | 11.06 | 71 | 72 | 72 | 71 | 71 | 0.55 |
| 21 | | | | | 11.28 | 109 | 107 | 106 | 106 | 106 | 1.30 |
| 18 | | | | | 17.75 | 170 | 172 | 173 | 171 | 173 | 1.30 |
| 21 | | | | | 11.28 | 109 | 107 | 106 | 106 | 106 | 1.30 |
| 19 | | | | | 10.15 | 73 | 74 | 74 | 75 | 75 | 0.84 |
| 20 | | | | | 8.38 | 92 | 91 | 88 | 91 | 95 | 2.51 |
| 22 | | | | | 13.94 | 129 | 133 | 130 | 130 | 130 | 1.52 |
| 29 | | | | | 13.11 | 76 | 78 | 78 | 78 | 79 | 1.10 |
| 51 | | | | | 4.34 | 41 | 44 | 41 | 41 | 41 | 1.34 |
| 52 | | | | | 5.49 | 43 | 44 | 46 | 46 | 43 | 1.52 |
| 54 | | | | | 6.57 | 54 | 53 | 56 | 54 | 54 | 1.10 |
| 55 | | | | | 6.14 | 53 | 51 | 55 | 54 | 51 | 1.79 |
| 58 | | | | | 12.47 | 78 | 76 | 75 | 76 | 76 | 1.10 |
| 59 | | | | | 7.94 | 51 | 52 | 49 | 51 | 52 | 1.22 |
| TOTAL: | **0** | **0** | **0** | **0** | **240.87** | | | | | AVG | 1.53 |
| | | | | | | | | | | MAX | 3.70 |
| | | | | | | | | | | MIN | 0.00 |

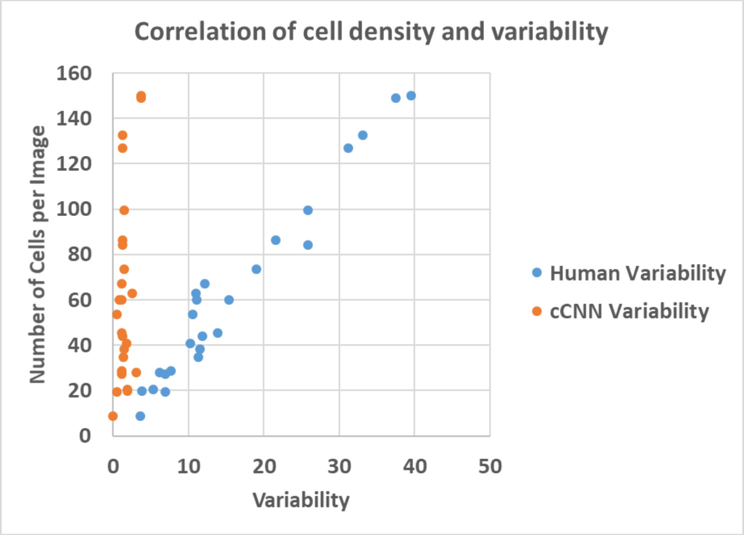

*Supplement Figure 8 – correlation between the number of cells found on microscope images, and human (blue) or software (orange) error.*

*Supplement Table 7 – Cell counts from repeat images (data recompiled from Supplement Table 6).*

| | im# | 1st Pass | 2nd Pass | STDEV |
|---|---|---|---|---|
| Person 1 | 1 | 23 | 28 | 3.5355 |
| | 2 | 37 | 35 | 1.4142 |
| | 21 | 99 | 98 | 0.7071 |
| | 18 | 164 | 151 | 9.1924 |
| | 56 | 194 | 192 | 1.4142 |
| | **im#** | **1st Pass** | **2nd Pass** | **STDEV** |
| Person 2 | 1 | 24 | 23 | 0.7071 |
| | 2 | 35 | 32 | 2.1213 |
| | 21 | 110 | 103 | 4.9497 |
| | 18 | 145 | 149 | 2.8284 |
| | 56 | 175 | 173 | 1.4142 |
| | **im#** | **1st Pass** | **2nd Pass** | **STDEV** |
| Person 3 | 1 | 16 | 15 | 0.7071 |
| | 2 | 21 | 19 | 1.4142 |
| | 21 | 53 | 56 | 2.1213 |
| | 18 | 89 | 85 | 2.8284 |
| | 56 | 106 | 102 | 2.8284 |
| | **im#** | **1st Pass** | **2nd Pass** | **STDEV** |
| Person 4 | 1 | 19 | 20 | 0.7071 |
| | 2 | 26 | 25 | 0.7071 |
| | 21 | 81 | 91 | 7.0711 |
| | 18 | 142 | 125 | 12.021 |
| | 56 | 136 | 143 | 4.9497 |
| | **im#** | **1st Pass** | **2nd Pass** | **STDEV** |
| Person 5 | 1 | 16 | 16 | 0 |
| | 2 | 21 | 21 | 0 |
| | 21 | 54 | 62 | 5.6569 |
| | 18 | 91 | 95 | 2.8284 |
| | 56 | 113 | 112 | 0.7071 |

*cCNN based cell counting algorithm* – this algorithm was based on a combination of methods used to preprocess microscope images and test the classification type neural network. The algorithm consisted of the following steps:

1. Load raw microscope image
2. Convert to binary
3. Bound all cell clusters >10 pixels in area with a bounding box.
4. Crop out sections of the raw image based on the coordinates acquired in step 3
5. Size normalize cropped images using the blended bumper method described in this study
6. Pass size normalized images to the advanced rotation network developed in this study
7. Record the prediction with the highest confidence value

**Example of microscope image used in this human performance study:**

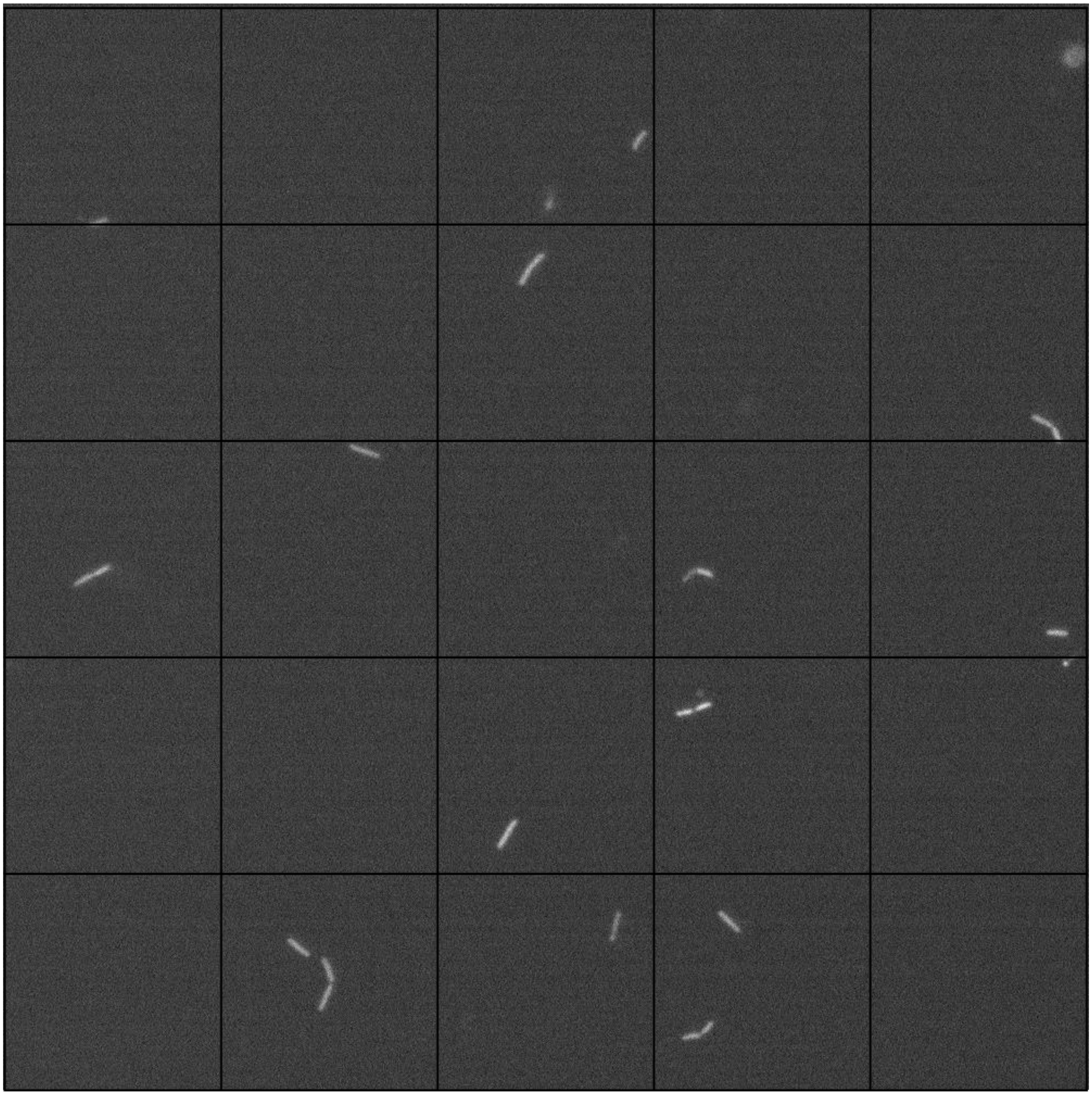

#### Section 13: image labeling GUI

The process of labeling microscope images is a tedious task. In order to streamline it, we developed a custom image labeling interface using Matlab. The app allows the user to load an image, segment it (binary thresholding), and present segments one at a time. The user can then review those segments and assign a class to the segment. This will generate a text file with the label for a specific segment. The app and code are made available at www.reuelgroup.org/resources.

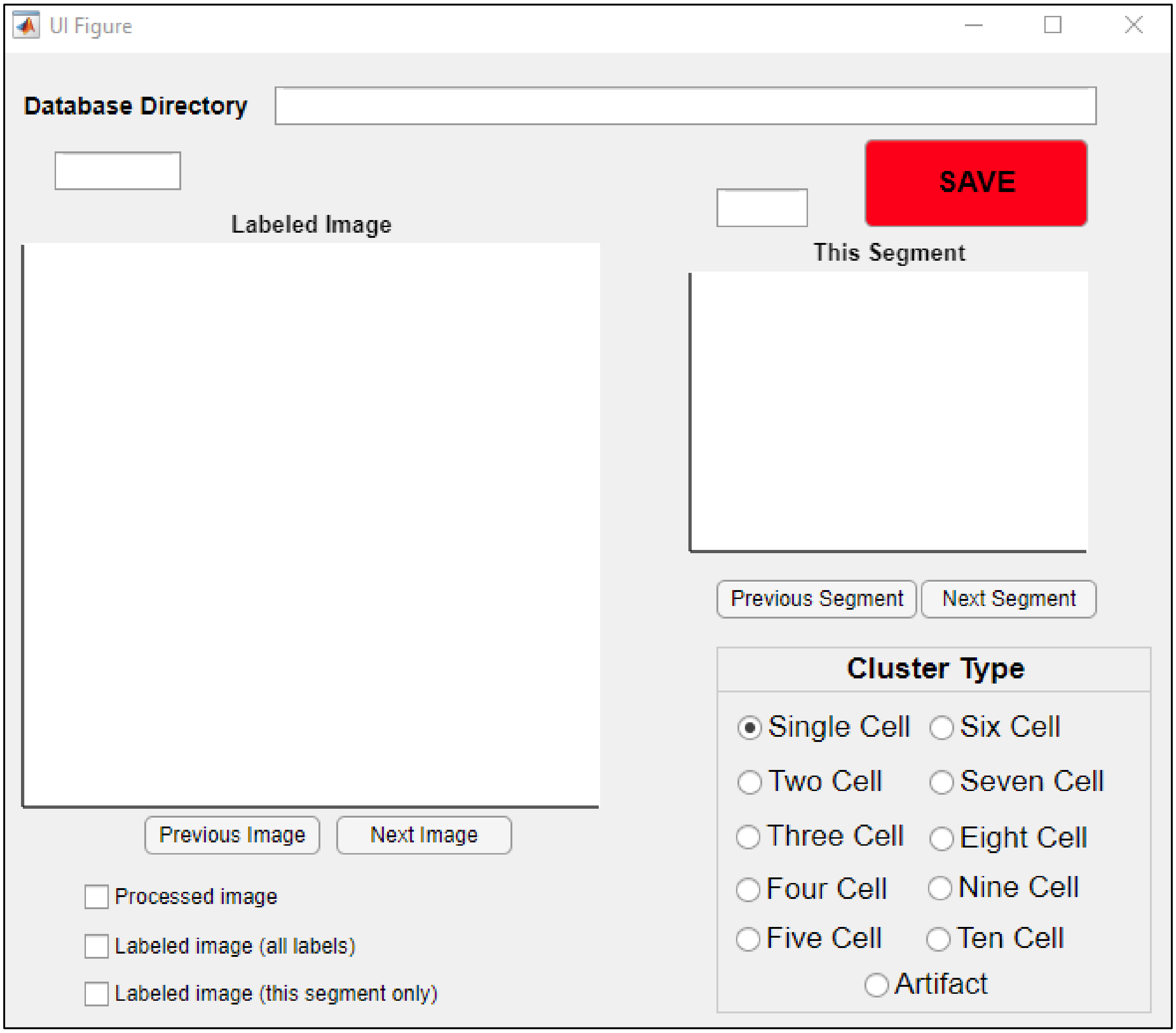

*Figure 1*
